# Application of High-Dimensional Statistics and Network based Visualization techniques on Arab Diabetes and Obesity data

**DOI:** 10.1101/151621

**Authors:** Raghvendra Mall, Reda Rawi, Ehsan Ullah, Khalid Kunji, Abdelkrim Khadir, Ali Tiss, Jehad Abubaker, Mohammed Dehbi, Halima Bensmail

**Author notes:** Equally contributing first authors (Raghvendra Mall, Reda Rawi). Corresponding author: Halima Bensmail, Qatar Computing Research Institute, Hamad Bin Khalifa University, Doha, Qatar.

## Abstract

**Background:** Obesity and its co-morbidities are characterized by a chronic low-grade inflammatory state, uncontrolled expression of metabolic measurements and dis-regulation of various forms of stress response. However, the contribution and correlation of inflammation, metabolism and stress responses to the disease are not fully elucidated. In this paper a cross-sectional case study was conducted on clinical data comprising 117 human male and female subjects with and without type 2 diabetes (T2D). Characteristics such as anthropometric, clinical and bio-chemical measurements were collected.

**Methods:** Association of these variables with T2D and BMI were assessed using penalized hierarchical linear and logistic regression. In particular, *elastic net, hdi* and *glinternet* were used as regularization models to distinguish between cases and controls. Differential network analysis using *closed-form* approach was performed to identify pairwise-interaction of variables that influence prediction of the phenotype.

**Results:** For the 117 participants, physical variables such as PBF, HDL and TBW had absolute coefficients 0.75, 0.65 and 0.34 using the *glinternet* approach, biochemical variables such as MIP, ROS and RANTES were identified as determinants of obesity with some interaction between inflammatory markers such as IL4, IL-6, MIP, CSF, Eotaxin and ROS. Diabetes was associated with a significant increase in thiobarbituric acid reactive substances (TBARS) which are considered as an index of endogenous lipid peroxidation and an increase in two inflammatory markers, MIP-1 and RANTES. Furthermore, we obtained 13 pairwise effects. The pairwise effects include pairs from and within physical, clinical and biochemical features, in particular metabolic, inflammatory, and oxidative stress markers.

**Conclusions:** We showcase that markers of oxidative stress (derived from lipid peroxidation) such as MIP-1 and RANTES participate in the pathogenesis of diseases such as diabetes and obesity in the Arab population.

## Introduction

Obesity has emerged as a major risk factor for the development of myriad chronic disorders that include insulin resistance (IR), type 2 diabetes (T2D), and metabolic syndrome [1, 2]. Moreover, poorly managed diabetes can lead to several micro- and macro-vascular complications such as heart failure, blindness, nephropathy, neuropathy and foot ulceration or amputation that may culminate in death [3, 4]. Of extreme concerns is the escalating rate by which obesity and diabetes are progressing across the world. According to the most recent estimations of the International Association for the Study of Obesity (www.iaso.org), the World Health Organization (www.who.org) and approximately 1.5 billion individuals worldwide were obese in 2015. The 2012 report of the International Diabetes Federation (www.idf.org) estimated the global number of diabetics to be about 371 million and it is projected to increase to about 552 million by 2030 if no proactive measures are promptly taken to control and prevent this epidemic disaster. Countries of the Gulf Cooperation Council (GCC) such as Saudi Arabia, Kuwait and Qatar have the highest prevalence of obesity and T2D in the world.

The pathophysiological mechanisms underlying these metabolic disorders involve a complex interplay between genetic, aging, behavioral, and environmental factors [5–7]. While genetic factors are key components in determining the susceptibility of individuals to weight gain and diabetes, they can be attenuated or exacerbated by a wide variety of modifiable factors involved in energy homeostasis, namely a sedentary lifestyle and behaviour, food intake, physical activity, smoking, and stress. Therefore, focus on populationbased public health interventions that target these modifiable factors associated with the development of these chronic diseases becomes an urgent task worldwide.

At the cellular level, obesity and diabetes are characterized by chronic low-grade inflammation and aberrant regulation of stress response in key metabolic organs such as adipose tissue, muscle and liver [8, 9]. The stress response; referred to as metabolic stress, is highly complex and includes persistent endoplasmic reticulum (ER)-mediated stress [10], enhanced oxidative stress [11], dysfunction of the mitochondria or defect in its biogenesis [12], hypoxia [13] and impairment of the host anti-stress defense system [14–17]. Recent evidence indicated that the uncontrolled inflammatory response and metabolic stress are highly integrated and they likely work in vicious cycles [9, 18, 19]. This represents one of the greatest challenges to identify therapeutic targets for the treatment and management of these metabolic disorders [9,20,21]. At the molecular level, the existence of such an environment leads to the activation of c-Jun NH2 terminal kinase (JNK) [22], and the inflammatory *κ*B kinase (IKK) [23]. Experimental evidence indicated clearly that JNK and IKK play a key role in the inhibition of the insulin receptor signaling cascade by virtue of their ability to phosphorylate and inactivate the insulin receptor substrate-1 (IRS-1), and thus, converting it to a poor substrate for the insulin receptor [18, 24].

In this case study, we carried out a multiplexing-based high throughput expression profiling of the inflammatory, metabolic and oxidative stress markers in human lean, overweight and obese subjects with and without T2D. A comprehensive statistical approach based on *elastic net* [25], *hdi* [26] and *glinternet* [27], was then undertaken to analyze the physical, clinical and biochemical data sets with the perspective to identifying the molecular signature specific for each group as well as the biological network of these signatures within and between the groups.

Our network based analysis using the **Closed-Form** approach [28] confirmed the close connection between obesity and T2D. In addition, it pointed to disease-responsive active modules and sub-clusters. Taken together, this approach should be helpful in the identification of novel biomarkers for the onset and progression of obesity, T2D, and associated diseases.

## Materials and Methods

### Study population

The study was conducted on 117 adult male and female human subjects with and without diabetes consisting of lean (Body mass index (BMI) = 18.5 – 24.9 kg/m^2^; n=20), overweight (BMI = 25 – 29.9 kg/m^2^; n=35) and obese (BMI = 30–40 kg/m^2^; n=62). Informed written consent was obtained from all subjects before their participation in the study, which was approved by the Review Board of Dasman Diabetes Institute and carried out in line with the guideline ethical declaration of Helsinki. Morbid obese (i.e. BMI *>* 40 kg/m^2^) and participants with prior major illness were excluded from the study. The physical characteristics of the participating subjects are shown in Tables 1 and 2.

**Table 1.**
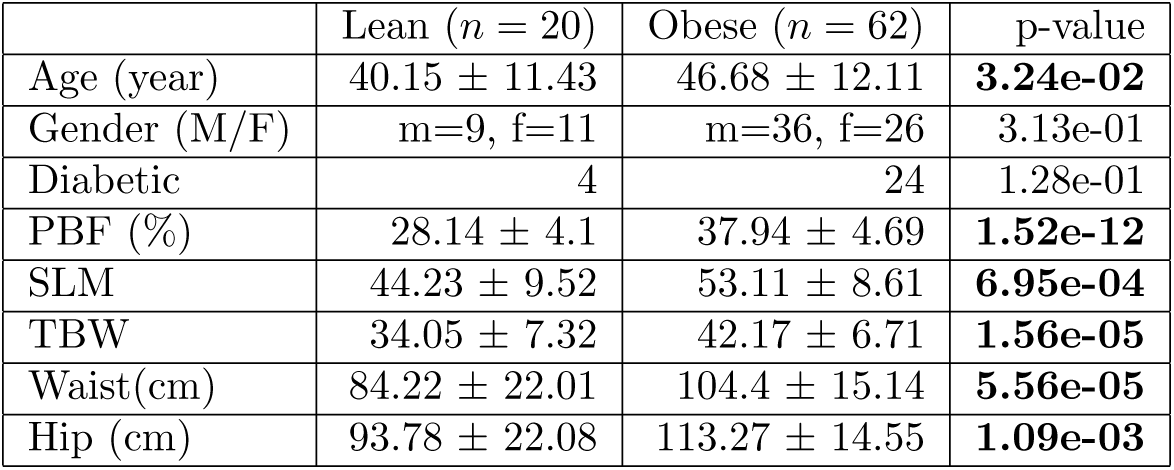
Physical characteristics of lean and obese subjects at baseline. Data are presented as mean ± SD. Here Percent body fat (PBF), Soft lean mass (SLM), Total body water (TBW).

**Table 2.** Physical characteristics of diabetic and non-diabetic subjects at baseline. Data are presented as mean ± SD. Here Body mass index (BMI), Percent body fat (PBF), Soft lean mass (SLM), Total body water (TBW).

| | Diabetic ($n = 36$) | Non-Diabetic ($n = 81$) | p-value |
| --- | --- | --- | --- |
| Age (year) | 52.08 ± 9.48 | 41.3 ± 11.68 | **3.56e-06** |
| Gender (M/F) | m=18, f=18 | m=48, f=33 | 3.56e-01 |
| BMI | 32.01 ± 4.08 | 29.74 ± 5.03 | **1.86e-02** |
| Weight (kg) | 87.33 ± 14.32 | 83.97 ± 15.92 | 2.19e-01 |
| Height (m) | 1.66 ± 0.08 | 1.68 ± 0.1 | 3.64e-01 |
| PBF (%) | 36.88 ± 5.56 | 33.37 ± 5.97 | **3.31e-03** |
| SLM | 50.07 ± 8.74 | 50.74 ± 9.49 | 5.87e-01 |
| TBW | 39.73 ± 6.71 | 39.92 ± 7.58 | 8.22e-01 |
| Waist (cm) | 100.89 ± 14.52 | 96.95 ± 18.6 | 2.63e-01 |
| Hip (cm) | 110.43 ± 12.29 | 104.5 ± 17.98 | **4.09e-02** |

### Anthropometric measurements, blood biochemistry and laboratory investigations

Anthropometric measurements were performed on all the participants. Whole-body composition was determined by dual-energy radiographic absorptiometry device (Lunar DPX, Lunar radiation, Madison, WI). Venous peripheral blood was collected from participants and used to prepare plasma and serum using standard methods. Glucose (GLU) and lipid profiles, including high-density lipoprotein (HDL) and low-density lipoprotein (LDL), were measured on the Siemens Dimension RXL chemistry analyzer (Diamond Diagnostics, Holliston, MA). Glycated haemoglobin (HbA1c) was determined using the VariantTM device (BioRad, Hercules, CA). Plasma levels of inflammatory and metabolic markers were measured using bead-based multiplexing technology using commercially available kits (BioRad, Hercules, CA). The panel of the inflammatory markers (##M500KCAF0Y) contains cytokines (IL-1*β*, IL-1ra, IL-4, IL-5, IL-6, IL-7, IL-8, IL-9, IL-10, IL-12 (p70), IL-13, IL-17, TNF-*α* and IFN-*γ*), chemokines (RANTES, IP-10, MCP1, MIP-1*α*, MIP-1*β*, Eotaxin) and growth factors (G-CSF and PDGF-BB,). The panel of metabolic markers (#171A7001M) contains 10 analytes consisting of (C-peptide, GIP, Ghrelin, Glucagon, GLP-1, Insulin, Leptin, PAI-1, Resistin and Visfatin). Median fluorescence intensities were collected on a Bioplex-200 system using Bioplex Manager software version 6 (BioRad, Hercules, CA). Lipid peroxidation was assessed by measuring plasma levels of malonaldehyde, using TBARs Assay Kit (Cayman Chemical Company, Ann Arbor, MI). Serum levels of ROS were determined using the OxiSelectTM ROS Assay Kit (Cell Biolabs Inc, San Diego, CA). Plasma/Serum levels of Paraoxonase 1 (PON1) were determined by using ELISA Kit (#ABIN414651 Life Technologies, Grand Island, New York, USA). All the above assays were carried out according to the instructions of the manufacturers.

### Missing value imputation

We identified that around 8% of the raw data are missing. Instead of removing the missing values we decided to approximate missing values using the well-known technique Multivariate Imputation by Chained Equations (MICE) implemented in R [29] package *mice* (https://cran.r-project.org/web/packages/mice/) [30].

### Data Analysis

Baseline statistical analysis of two groups in each dataset were calculated using R. Statistics for all the variables in the study are reported as means ± standard deviation (SD) unless otherwise stated. The R implementation of the Anderson-Darling test in the *nortest* package (https://cran.r-project.org/web/packages/nortest/) [31] was used to test for normality of all the variables. If a variable is not normally distributed in both groups, the MannWhitney test was used to determine significance of the difference in means between the groups. For a normally distributed variable in both groups, the Student’s t-test was used to determine significance of difference in means between groups. In this case, the F-test was used to compare variance of the variable in the groups. A p-value lower than 0.05 indicates a statistically significant difference between the groups.

### Regularization models

We utilize a linear regression model with *n* observations and *p* explanatory variables (features)

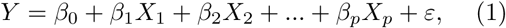

where *Y* = (*y*_1_, …, *y_n_*)*^t^* is the response, *ε* = (*ε*_1_, …, *ε_n_*)*^t^* ~ *N*(0, *σ*^2^*I_n_*) is the noise vector; *X_j_* represents the *j^th^* predictor and *β* = (*β*_1_, …, *β_p_*)*^t^* is the vector of parameters of interest to be estimated; each *β_j_*, *j* = 1, …, *p* represents the association between the variable *X_j_* (feature) and the response *Y.* The greater the absolute value of *β,* the stronger is the effect of the corresponding feature.

### Elastic Net

The LASSO coefficients, 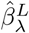, minimize the quantity

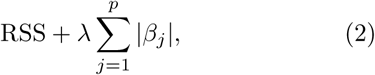

with RSS as the *residual sum of squares* and *λ* as the tuning parameter. The LASSO technique penalizes hereby the regression coefficients using an *L*_1_ norm. The *L*_1_ penalty has the effect of forcing some of the coefficient to be exactly equal to zero when the tuning parameter *λ* is sufficiently large. Hence, the LASSO estimates the coefficients and performs variable selection at the same time [32].

The *elastic net* regularization regression method introduced in [33] combines the *L*_1_ and *L*_2_ penalties and overcomes among others the following limitations of the classical LASSO:

- In *p* > *n* cases, the LASSO selects maximum n variables when converging, which is limiting characteristic of a variable selection method.
- LASSO selects only one variable from a group of variables that have high pairwise correlations

The coefficients from the *elastic net* are formulated as follows:

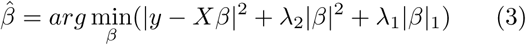

We used R package *glmnet* (https://cran.r-project.org/web/packages/glmnet/) [34] to calculate the 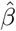 coefficients. We performed 10-fold cross validation while training the *elastic net* model.

### High-dimensional inference

In the case of *p* > *n* it is not possible to use the covariance test without specifying an estimate of the error standard deviation i.e.∑^2^. Meinshausen et al. introduced in [35] an approach where the data is split into two groups LASSO regularization, in particular *elastic net* 10-fold cross validation, is applied on one group where-after the variables selected by LASSO are used as predictors to obtain p-values from an ordinary least squared regression on the other group. We used R package *hdi* (https://cran.r-project.org/web/packages/hdi/) [36] to calculate the p-values.

### Glinternet

In order to study the interaction effects of features, we applied Lim and Hastie’s approach *glinternet* (https://cran.r-project.org/web/packages/glinternet/) [37]. This method learns pairwise interactions in a regression model that satisfies hierarchy constraints. Further and to the best of our knowledge, this is the only approach that allows a mixture of categorical and continuous values which is the case with our data.

We used R package *glinternet* to generate the main and interaction coefficients. We performed 10-fold cross validation when training a *glinternet* interaction model.

### Network Based Analysis

We have applied several statistical methods to identify variables or variable interactions which help to distinguish control from patient for diabetes and lean from obese w.r.t. BMI as already introduced. Here, we perform network based analysis to identify differential variables and their interaction for the same set of problems.

### Network Construction

We first construct networks for interactions between the variables for the two groups in datasets 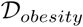 and 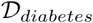. Here 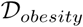 comprises all the people who are either obese or lean and 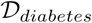 consists of all the people who are either diabetic or non-diabetic. Each variable is considered as a node in the network and let 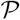 represent the set of all the variables/nodes. An edge between two nodes *i* and *j* is induced by calculating the mutual information (MI) between two variables. It is well known from information theory that MI is a measure of mutual dependence between two random variables. Higher values of MI indicate that the variables are dependent while values ≈ 0 represent that the variables are mutually independent i.e. change in one variable does not effect the other. By performing this operation, we obtain mutual information 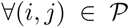 thereby resulting in a full interaction graph between the variables for a particular case.

To ensure the robustness of the generated networks we apply a nonparametric bootstrap procedure [38]. This provides for each node a minimum value of MI which is necessary for its edge to be included in the final network. As a result of this procedure we remove all non-significant edges from the network making it sparse. We then convert these networks into topological overlap graphs [28, 39] i.e. the edge weights quantify the topological overlap (TO) between a pair of nodes by taking into account the local neighbourhood structure around those nodes [40]. This results in symmetric, undirected and weighted networks that are used for differential subnetwork analysis as indicated in [28]. Finally, we remove all self-loops from the topological network along with removal of any isolated node i.e. nodes with no connections. By performing this operation we reduce the size of the interaction networks as showcased in the results section.

### Differential Network Analysis

We utilize the **Closed-Form** differential subnetwork analysis technique proposed in [28] to identify statistically significant subgraphs when performing paired network comparison i.e. when comparing variable interaction network (topological graphs) for lean with obese case and control with patient case for diabetes. We briefly explain the Generalized Hamming Distance used to estimate the distance between two graphs. Given two topological networks 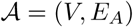 and 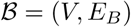 where *V* represents the set of nodes i.e. 1, …, *N* and *E_i_* represents the edges in the *i^th^* network. The hamming distance between *A* and *B* is given by 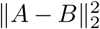 which represents the Frobenius norm of the difference between *A* and *B* graphs. The Generalized Hamming Distance (GHD) is defined as:

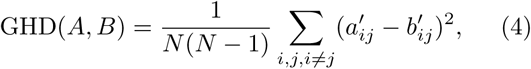

where 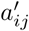 and 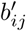 are mean centered edge-weights defined as:

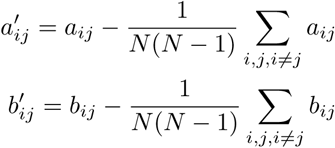

Ruan et al. proposed the method differential Generalized Hamming Distance (dGHD) to obtain closed-form p-values for the null hypothesis that *A* and *B* are independent [39]. They efficiently calculate the p-value and circumvent expensive permutation processes by assuming asymptotic normality. This can be represented as:

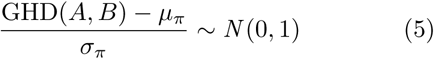

Here *μ_π_* is the asymptotic value of the mean GHD and *σ_π_* is the asymptotic value of the standard deviation of the GHD for permutations of *A* w.r.t. *B.* In order to estimate the *μ_π_* and *σ_π_* values we define:

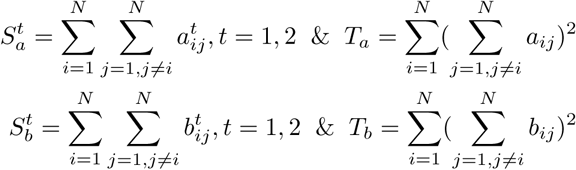

Here 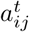 and 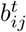 are the edge weights with the power *t.* Furthermore, we require the following terms:

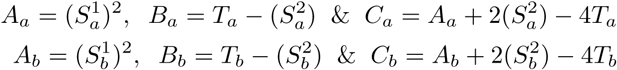

The notion of differential subnetworks is based on the idea that when comparing two networks only a subset of edges would present altered interaction. The goal is to identify those set of nodes associated with such a subset of edges. For this subset *V\** there is no sufficient evidence to reject the null hypothesis that the corresponding subnetworks *A\**(*V\**, *E_A*_)* and *B\**(*V\**, *E_B*_*) are statistically independent. We utilized the more advanced **Closed-Form** algorithm [28], which is computationally cheaper and detects fewer false positives w.r.t. the dGHD [39] technique, for identifying the differential subnetworks.

## Results

We removed physical characteristics namely height and weight while performing the analysis for obesity. Similarly, we removed clinical characteristics namely blood glucose (GLU) and HbA1c when analysing diabetes. This is because these traits are often used to measure obesity and diabetes respectively (hence they act as confounding variables when performing the analysis for obesity and diabetes).

### Baseline Characteristics of Study Population

Physical characteristics of datasets 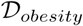 and 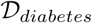 are summarized in Table 1 and 2 respectively. Age, percent body fat (PBF), soft lean mass (SLM), total body weight (TBW), waist and hip size were found significantly higher (p-value: 3.24e-02, 5.51e-10, 1.52e-12, 6.95e-04, 1.56e-05, 5.56e-05 and 1.09e-03 respectively) in the obese compared to lean subjects as expected. Age, BMI, PBF, and hip size were found significantly higher (p-value: 3.56e-06, 1.86e-02, 3.31e-03 and 4.09e-02 respectively) in the diabetic subjects compared to non-diabetic subjects.

Clinical characteristics of datasets 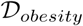 and 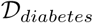 are summarized in Table 3 and 4 respectively. Obese subjects have significantly higher levels of triglycerides (TGL) compared to lean subjects (p-value: 1.25e-02).

**Table 3.** Clinical characteristics of lean and obese subjects at baseline. In our study we have not considered the overweight case to have a clear distinction between lean and obese cases. Data are presented as mean ± SD. Here Cholesterol (Chol), High density lipoprotein (HDL), Low density lipoprotein (LDP), and Triglycerides (TGL).

| | Lean ($n = 20$) | Obese ($n = 62$) | p-value |
| --- | --- | --- | --- |
| Chol (mmol/l) | $4.96 \pm 0.8$ | $5.18 \pm 1.05$ | 4.76e-01 |
| HDL (mmol/l) | $1.26 \pm 0.33$ | $1.18 \pm 0.36$ | 3.89e-01 |
| LDL (mmol/l) | $3.21 \pm 0.76$ | $3.23 \pm 1.33$ | 9.12e-01 |
| TGL (mmol/l) | $1.08 \pm 0.53$ | $2.11 \pm 3.03$ | **1.25e-02** |

**Table 4.** Clinical characteristics of diabetic and non-diabetic subjects at baseline. Data are presented as mean ± SD. Here Cholesterol (Chol), High density lipoprotein (HDL), Low density lipoprotein (LDP), and Triglycerides (TGL).

| | Diabetic ($n = 36$) | Non-Diabetic ($n = 81$) | p-value |
| --- | --- | --- | --- |
| Chol (mmol/l) | $5.05 \pm 1.18$ | $5.19 \pm 0.91$ | 3.77e-01 |
| HDL (mmol/l) | $1.27 \pm 0.44$ | $1.21 \pm 0.38$ | 4.9e-01 |
| LDL (mmol/l) | $3.05 \pm 1.58$ | $3.32 \pm 0.9$ | 3.54e-01 |
| TGL (mmol/l) | $2.48 \pm 3.87$ | $1.36 \pm 0.82$ | 9.22e-02 |

Metabolic profiles of datasets 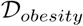 and 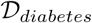 are summarized in Table 5 and 6 respectively. Levels of insulin, leptin, Plasminogen activation inhibitor (PAI-1), Interleukin 13 (IL-13), Interferon-gamma-inducible protein-10 (IP-10), Reactive oxygen species (ROS) and Thiobarbituric acid reactive substances (TBARS) are found significantly higher in obese compared to lean subjects (p-value: 4.02e-04, 4.08e-03, 4.52e-02, 1.68e-02, 7.64e-03, 5.69e-03 and 1.04e-02 respectively). Levels of MIP-1*α* and TBARS are found significantly higher in diabetic subjects compared to non-diabetic subjects (p-value: 3.86e-02 and 5.96e-04 respectively).

**Table 5.** Biochemical characteristics of lean and obese subjects at baseline. Data are presented as mean ± SD. Here Gastric inhibitory peptide (GIP), Glucagon like peptide-1 (GLP-1), Granulocyte colony stimulating factor (G-CSF), Interleukin (IL), Interleukin-1 receptor agonist (IL-1ra), Interferon-gamme (IFN-*γ*), Interferon-gamma-inducible protein-10 (IP-10), Monocyte chemoattractant protein-1 (MCP-1), Macrophage inflammatory protein-1*α* (MIP-1*α*), Macrophage inflammatory protein-1*β* (MIP-1*β*), Platelet-derived growth factor-bb (PDGF-bb), Tumor necrosis factor-*α* (TNF-*α*), Paraoxonase-1 (PON-1), Reactive oxygen species (ROS), Thiobarbituric Acid Reactive Substances (TBARS).

| | Lean ($n = 20$) | Obese ($n = 62$) | p-value |
| --- | --- | --- | --- |
| Metabolic markers |  |  |  |
| C-peptide (ng/ml) | 2437.75 ± 733.17 | 2864.74 ± 1251.84 | 6.67e-02 |
| GIP (pg/ml) | 151.59 ± 69.09 | 162.76 ± 86.5 | 6.01e-01 |
| Ghrelin (pg/ml) | 151 ± 82.69 | 145.39 ± 108.66 | 8.33e-01 |
| Glucagon (ng/ml) | 673.85 ± 93.28 | 684.43 ± 137.14 | 4.34e-01 |
| GLP-1 (ng/ml) | 2541.66 ± 909.12 | 2551.85 ± 1341.7 | 9.75e-01 |
| Insulin (ng/ml) | 2421.87 ± 1035.68 | 4015.97 ± 2864.78 | **4.02e-04** |
| Leptin (ng/ml) | 4955.66 ± 3048.97 | 8167.55 ± 4527.31 | **4.08e-03** |
| PAI-1 (ng/ml) | 3063.25 ± 1590.61 | 3704.57 ± 1388.07 | **4.52e-02** |
| Resistin (ng/ml) | 1208.4 ± 515.89 | 968.31 ± 462.72 | 5.33e-02 |
| Visfatin (ng/ml) | 9139.89 ± 5148.53 | 9225.14 ± 7737.6 | 9.63e-01 |
| Inflammatory markers |  |  |  |
| IL-1β (pg/ml) | 1.13 ± 0.52 | 1.32 ± 0.88 | 2.49e-01 |
| IL-1ra (pg/ml) | 95.59 ± 41.84 | 91.21 ± 46.44 | 7.08e-01 |
| IL-4 (pg/ml) | 2.17 ± 1.03 | 1.95 ± 0.98 | 3.93e-01 |
| IL-5 (pg/ml) | 2.18 ± 0.78 | 2.41 ± 1.14 | 4.05e-01 |
| IL-6 (pg/ml) | 5.13 ± 2.1 | 4.9 ± 2.07 | 6.63e-01 |
| IL-7 (pg/ml) | 5.15 ± 1.69 | 5.36 ± 2.12 | 6.93e-01 |
| IL-8 (pg/ml) | 5.68 ± 1.37 | 6.15 ± 3.65 | 4e-01 |
| IL-9 (pg/ml) | 13.9 ± 10.74 | 12.7 ± 9.6 | 6.39e-01 |
| IL-10 (pg/ml) | 1.61 ± 0.96 | 2.07 ± 2.29 | 2.02e-01 |
| IL-12 (p70) (pg/ml) | 7.42 ± 5.08 | 9.52 ± 5.79 | 1.52e-01 |
| IL-13 (pg/ml) | 2.48 ± 1.12 | 3.71 ± 3.46 | **1.68e-02** |
| IL-17 (pg/ml) | 12.61 ± 12.08 | 11.3 ± 10.73 | 6.48e-01 |
| Eotaxin (pg/ml) | 29.6 ± 20.2 | 39.11 ± 38.79 | 1.6e-01 |
| G-CSF (pg/ml) | 40.12 ± 15.23 | 42.46 ± 14.09 | 5.27e-01 |
| IFN-γ (pg/ml) | 45.16 ± 22.23 | 44.24 ± 26.24 | 8.88e-01 |
| IP-10 (pg/ml) | 393.99 ± 236.34 | 592.28 ± 378.7 | **7.64e-03** |
| MCP-1 (pg/ml) | 9.4 ± 2.52 | 10.32 ± 4.91 | 2.74e-01 |
| MIP-1α (pg/ml) | 8.66 ± 16.66 | 6.05 ± 9.25 | 5.11e-01 |
| PDGF-BB (pg/ml) | 531 ± 672.13 | 492.41 ± 589.44 | 8.06e-01 |
| MIP-1β (pg/ml) | 22.36 ± 6.6 | 27.07 ± 27.16 | 2.13e-01 |
| RANTES (pg/ml) | 1298.49 ± 635.18 | 1596.9 ± 751.28 | 1.14e-01 |
| TNF-α (pg/ml) | 25.19 ± 9.89 | 26.91 ± 11.79 | 5.57e-01 |
| Oxidative stress markers |  |  |  |
| PON (U) | 0.38 ± 0.11 | 0.37 ± 0.1 | 9.44e-01 |
| ROS (M) | 1426.07 ± 251.89 | 1608.57 ± 168.97 | **5.69e-03** |
| TBARS (μM) | 1.29 ± 0.6 | 1.77 ± 0.74 | **1.04e-02** |

**Table 6.** Biochemical characteristics of diabetic and non-diabetic subjects at baseline. Data are presented as mean ± SD. Here Gastric inhibitory peptide (GIP), Glucagon like peptide-1 (GLP-1), Granulocyte colony stimulating factor (G-CSF), Interleukin (IL), Interleukin-1 receptor agonist (IL-1ra), Interferon-gamme (IFN-*γ*), Interferon-gamma-inducible protein-10 (IP-10), Monocyte chemoattractant protein-1 (MCP-1), Macrophage inflammatory protein-1*α* (MIP-1*α*), Macrophage inflammatory protein-1*β* (MIP-1*β*), Platelet-derived growth factor-bb (PDGF-bb), Tumor necrosis factor-*α* (TNF-*α*), Paraoxonase-1 (PON-1), Reactive oxygen species (ROS), Thiobarbituric Acid Reactive Substances (TBARS).

| | Diabetic ($n = 36$) | Non-Diabetic ($n = 81$) | p-value |
| --- | --- | --- | --- |
| Metabolic markers |  |  |  |
| C-peptide (ng/ml) | 2482.96 ± 975.2 | 2761.7 ± 1182.23 | 2.18e-01 |
| GIP (pg/ml) | 160.72 ± 79.25 | 150.51 ± 87.52 | 5.5e-01 |
| Ghrelin (pg/ml) | 145.44 ± 94.87 | 146.3 ± 99.62 | 9.65e-01 |
| Glucagon (ng/ml) | 668.72 ± 108.61 | 669.8 ± 135.61 | 7.79e-01 |
| GLP-1 (ng/ml) | 2412.05 ± 1018.62 | 2596.7 ± 1297.95 | 4.51e-01 |
| Insulin (ng/ml) | 4136.91 ± 3338.54 | 2990.7 ± 1830.14 | 5.94e-02 |
| Leptin (ng/ml) | 7158.54 ± 4457.82 | 6702.58 ± 3893.55 | 5.77e-01 |
| PAI-1 (ng/ml) | 3576.96 ± 1254.45 | 3290.12 ± 1514.19 | 3.22e-01 |
| Resistin (ng/ml) | 1043.63 ± 463.53 | 1028.91 ± 456.37 | 8.73e-01 |
| Visfatin (ng/ml) | 8316.41 ± 4961.89 | 9470.67 ± 7847.87 | 3.39e-01 |
| Inflammatory markers |  |  |  |
| IL-1β (pg/ml) | 1.2 ± 0.83 | 1.22 ± 0.68 | 8.95e-01 |
| IL-1ra (pg/ml) | 93.73 ± 42.88 | 91.92 ± 43.16 | 8.34e-01 |
| IL-4 (pg/ml) | 1.84 ± 0.83 | 2.07 ± 1.07 | 2.54e-01 |
| IL-5 (pg/ml) | 2.16 ± 0.72 | 2.41 ± 1.12 | 1.57e-01 |
| IL-6 (pg/ml) | 4.7 ± 1.58 | 4.91 ± 2.09 | 5.95e-01 |
| IL-7 (pg/ml) | 4.91 ± 1.85 | 5.31 ± 1.88 | 2.84e-01 |
| IL-8 (pg/ml) | 6.4 ± 4.52 | 5.63 ± 1.67 | 3.26e-01 |
| IL-9 (pg/ml) | 12.21 ± 8.2 | 12.97 ± 10.21 | 6.94e-01 |
| IL-10 (pg/ml) | 1.54 ± 1.09 | 1.92 ± 2.05 | 1.92e-01 |
| IL-12 (p70) (pg/ml) | 7.88 ± 5.12 | 9 ± 5.16 | 2.83e-01 |
| IL-13 (pg/ml) | 3.15 ± 1.82 | 3.5 ± 3.15 | 4.53e-01 |
| IL-17 (pg/ml) | 8.77 ± 8.9 | 12.91 ± 11.52 | 5.81e-02 |
| Eotaxin (pg/ml) | 31.6 ± 19.46 | 39.41 ± 35.38 | 1.28e-01 |
| G-CSF (pg/ml) | 38.42 ± 12.87 | 42.59 ± 14.11 | 1.2e-01 |
| IFN-γ (pg/ml) | 40.57 ± 17.46 | 45.75 ± 25.55 | 2.05e-01 |
| IP-10 (pg/ml) | 570.47 ± 494.21 | 467.13 ± 218.56 | 2.36e-01 |
| MCP-1 (pg/ml) | 10.16 ± 4.86 | 9.84 ± 3.66 | 7.24e-01 |
| MIP-1α (pg/ml) | 8.76 ± 11.55 | 4.52 ± 9.45 | **3.86e-02** |
| PDGF-BB (ng/ml) | 464.06 ± 568.28 | 526.34 ± 641.18 | 6.17e-01 |
| MIP-1β (pg/ml) | 21.18 ± 8.62 | 26.08 ± 23.56 | 1.04e-01 |
| RANTES (ng/ml) | 1258.59 ± 593.56 | 1464.76 ± 744.95 | 1.46e-01 |
| TNF-α (pg/ml) | 26.43 ± 10.83 | 26.85 ± 11.99 | 8.57e-01 |
| Oxidative stress markers |  |  |  |
| PON (U) | 0.37 ± 0.1 | 0.36 ± 0.1 | 7.03e-01 |
| ROS (M) | 1542.61 ± 189.22 | 1546.04 ± 194.95 | 9.3e-01 |
| TBARS (μM) | 1.94 ± 0.81 | 1.4 ± 0.54 | **5.96e-04** |

### Regularisation models

#### BMI

We studied the effects of physical, clinical and biochemical features w.r.t. to lean and obese cases by applying *elastic net, hdi* and *glinternet.* We distinguish hereby between lean and obese cases. Throughout this section we will only list coefficients that are non-zero and p-values below a significance threshold of 0.05.

In Table 7, we list the coefficients and p-values obtained for different features when by applying *elastic net* and *hdi.* The features are sorted according to their effect strength (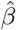 absolute values). The features with the highest *elastic net* coefficients include height, HDL, PBF, and TBW with 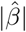 equal to 0.75, 0.44, and 0.16 respectively. The multi-sample splitting method implemented in *hdi* yielded two features as highly significant to distinguish between lean and obese cases. In particular, these characteristics are PBF and TBW with corrected p-values of 1.49e-09 and 6.29e-06.

**Table 7.** *Elastic net* and *hdi* results for BMI study

| | *elastic net* coefficient ($\hat{\beta}$) | *hdi* significant p-value |
| --- | --- | --- |
| HDL | -0.75 |  |
| PBF | 0.44 | 1.49e-09 |
| TBW | 0.16 | 6.29e-06 |
| SLM | 0.06 |  |
| Age | 0.02 |  |
| Waist | 0.01 |  |
| MIP-1β | 4.54e-03 | |
| MIP-1α | -3.79e-03 | |
| ROS | 1.41e-03 |  |
| RANTES | 5.78e-04 |  |
| Insulin | 7.12e-05 |  |

In Table 8 we summarized the single and pairwise coefficients obtained by applying the *glinternet* approach. Interestingly, we observed several main and pairwise non-zero coefficients. The main effects comprised the expected physical characteristics PBF, HDL and TBW with coefficients 0.75, -0.65, and 0.34. We also obtained a coefficient for the inflammatory marker RANTES, in particular with a coefficient 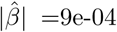. Next to the main effects, we obtained 13 interesting pairwise effects that describe the best model that distinguishes between lean and obese cases. The non-zero pairwise coefficients represent pairs of markers of different types, such as physical, clinical, as well as metabolic, inflammatory, and oxidative stress markers.

**Table 8.** *Glinternet* results for BMI study

| 1<sup>st</sup> feature | 2<sup>nd</sup> feature | *glinternet* coefficient ($\hat{\beta}$) |
| --- | --- | --- |
| PBF |  | 0.75 |
| HDL |  | -0.65 |
| TBW |  | 0.34 |
| RANTES |  | 9e-04 |
| Age | PON | 1.61e-02 |
| IL-6 | G-CSF | -7.55e-03 |
| IL-4 | MIP-1α | -6.08e-03 |
| GLU | Eotaxin | -5.33e-03 |
| SLM | TBW | -2.06e-03 |
| TBW | MIP-1β | -1.2e-03 |
| HDL | PAI-1 | 4.52e-04 |
| TBW | IL-1ra | 6.34e-05 |
| PBF | ROS | -5.03e-05 |
| Age | Glucagon | 3.39e-05 |
| GIP | Glucagon | 2.88e-06 |
| Age | Resistin | 4.01e-07 |
| Insulin | RANTES | -1.87e-07 |

#### Diabetes

In this subsection, we report the effects of physical, clinical and biochemical features on diabetes applying the same set of regularization methods. In Table 9, we listed the results obtained using *elastic net* and *hdi.* Unlike the BMI case, *elastic net* provided fewer features with non-zero coefficients. In particular, we observed the highest coefficient for the oxidative stress marker TBARS with 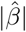 equal to 0.3. Further, we obtained coefficients for the physical marker age and PBF and the clinical marker TGL. The multi-sample splitting method *hdi* did not provide significant p-values to distinguish between diabetic and control cases.

**Table 9.** *Elastic net* and *hdi* results for Diabetes study

| | *elastic net* coefficient ($\hat{\beta}$) | *hdi* significant p-value |
| --- | --- | --- |
| TBARS | 0.3 |  |
| Age | 0.03 |  |
| TGL | 0.02 |  |
| PBF | 0.02 |  |

In Table 10 we listed the single and pairwise coefficients for the diabetes study obtained using *glinternet.* Interestingly, we observed many main and pairwise non-zero coefficients. The main effects include the oxidative stress marker TBARS, the clinical marker TGL, the physical characteristic age, and two inflammatory markers MIP-1*β* and RANTES. Furthermore, we obtained 13 pairwise effects with coefficients ranging from -5.03e-05 to 1.61e-02. The pairwise effects include pairs from and within physical, clinical and all three biochemical feature classes, in particular metabolic, inflammatory, and oxidative stress markers.

**Table 10.**
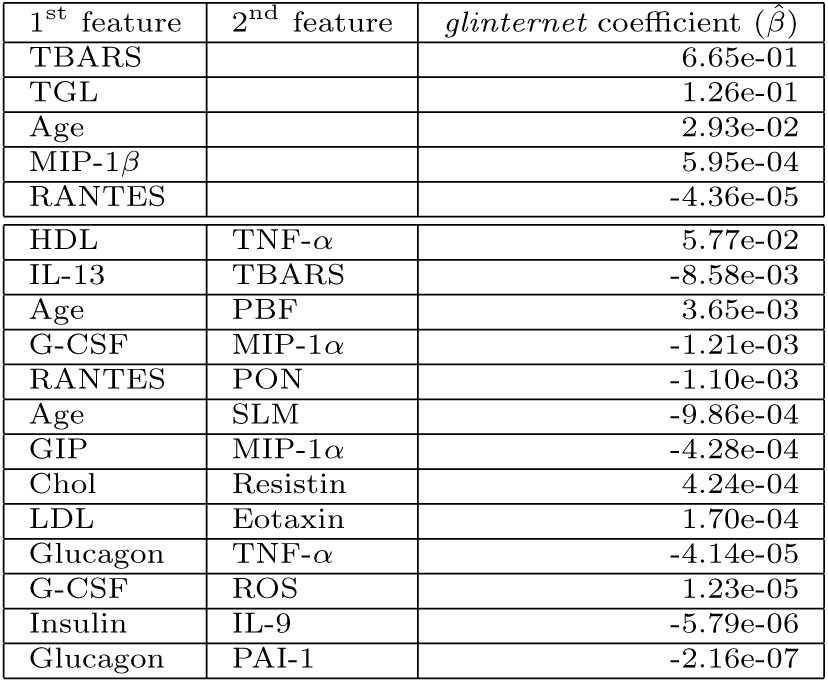
*Glinternet* results for Diabetes study

| 1<sup>st</sup> feature | 2<sup>nd</sup> feature | *glinternet* coefficient ($\hat{\beta}$) |
| --- | --- | --- |
| TBARS |  | 6.65e-01 |
| TGL |  | 1.26e-01 |
| Age |  | 2.93e-02 |
| MIP-1β | | 5.95e-04 |
| RANTES |  | -4.36e-05 |
| HDL | TNF-α | 5.77e-02 |
| IL-13 | TBARS | -8.58e-03 |
| Age | PBF | 3.65e-03 |
| G-CSF | MIP-1α | -1.21e-03 |
| RANTES | PON | -1.10e-03 |
| Age | SLM | -9.86e-04 |
| GIP | MIP-1α | -4.28e-04 |
| Chol | Resistin | 4.24e-04 |
| LDL | Eotaxin | 1.70e-04 |
| Glucagon | TNF-α | -4.14e-05 |
| G-CSF | ROS | 1.23e-05 |
| Insulin | IL-9 | -5.79e-06 |
| Glucagon | PAI-1 | -2.16e-07 |

### Differential Network Analysis

#### BMI

In Figure 1 we summarise significant mutual information (MI) values of all variable pairs for the dataset 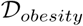 as heat maps (see Methods). The heat maps were generated using *heatmap.2* function in R package *gplots* (https://cran.r-project.org/web/packages/gplots/) [41]. In the lean subjects, as shown in Figure 1A, we observe two predominant clusters where the paired variables have high mutual dependence whereas in the obese case depicted in Figure 1B we see several clusters with relatively lower mutual dependence between the variables within the clusters. To highlight the subtle differences between the lean and obese cases we utilised the Closed-Form technique.

**Fig 1.**
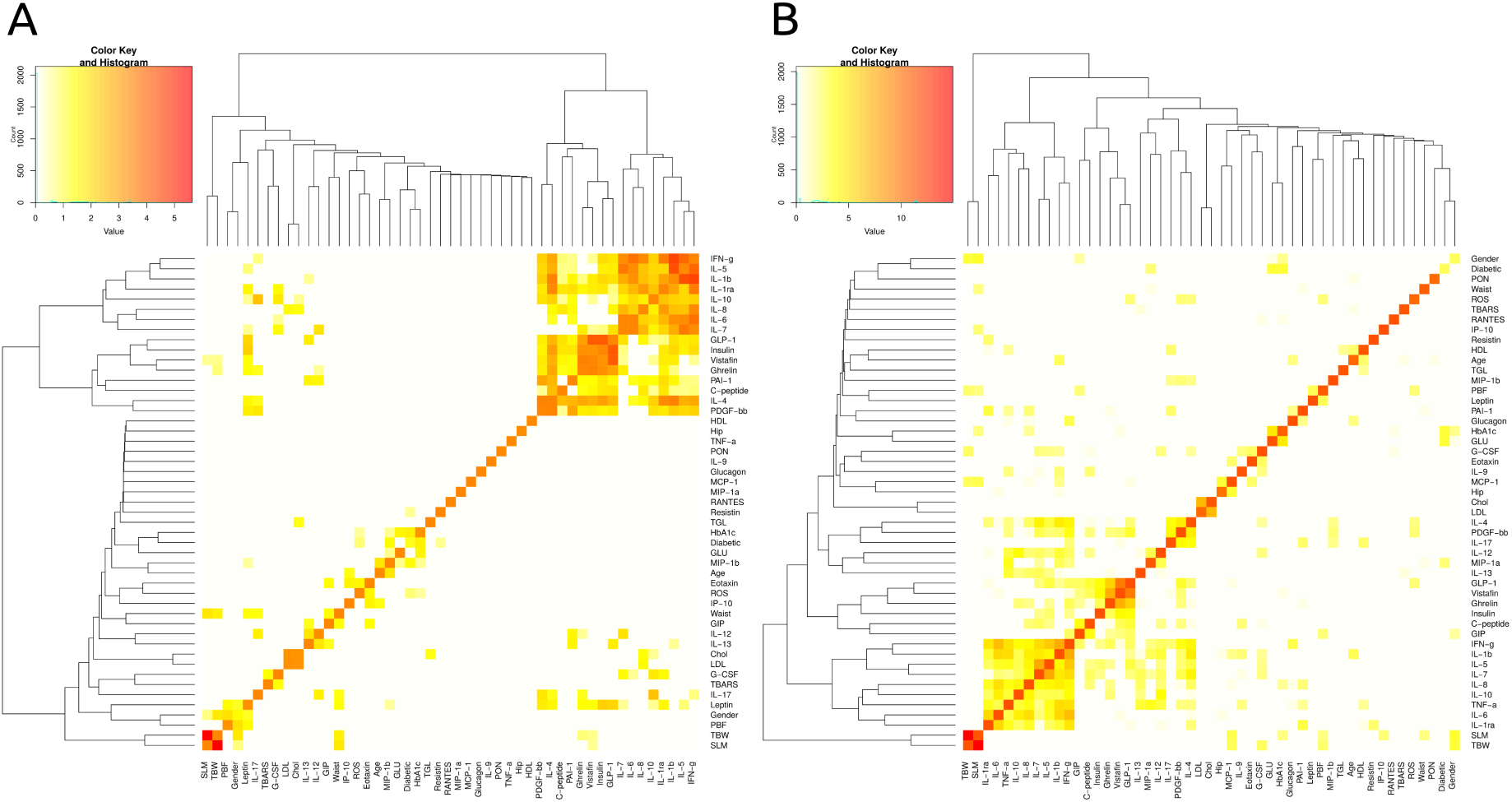
Mutual information heat map for the 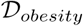 data set. MI based heat map of variables representing lean cases (A) and obese cases (B).

First, we show in Figure 2 the mutual dependance networks for lean *G_lean_* (Figure 2A) and obese *G_obese_* cases (Figure 2B). The *G_lean_* network comprises 40 nodes with 716 edges whereas *G_obese_* consists of 49 node and 1272 edges. We used the Louvain method [42] for the task of identifying communities [43–45] in all the networks that we built. We identified five clusters in both networks using the Louvain method.

**Fig 2.**
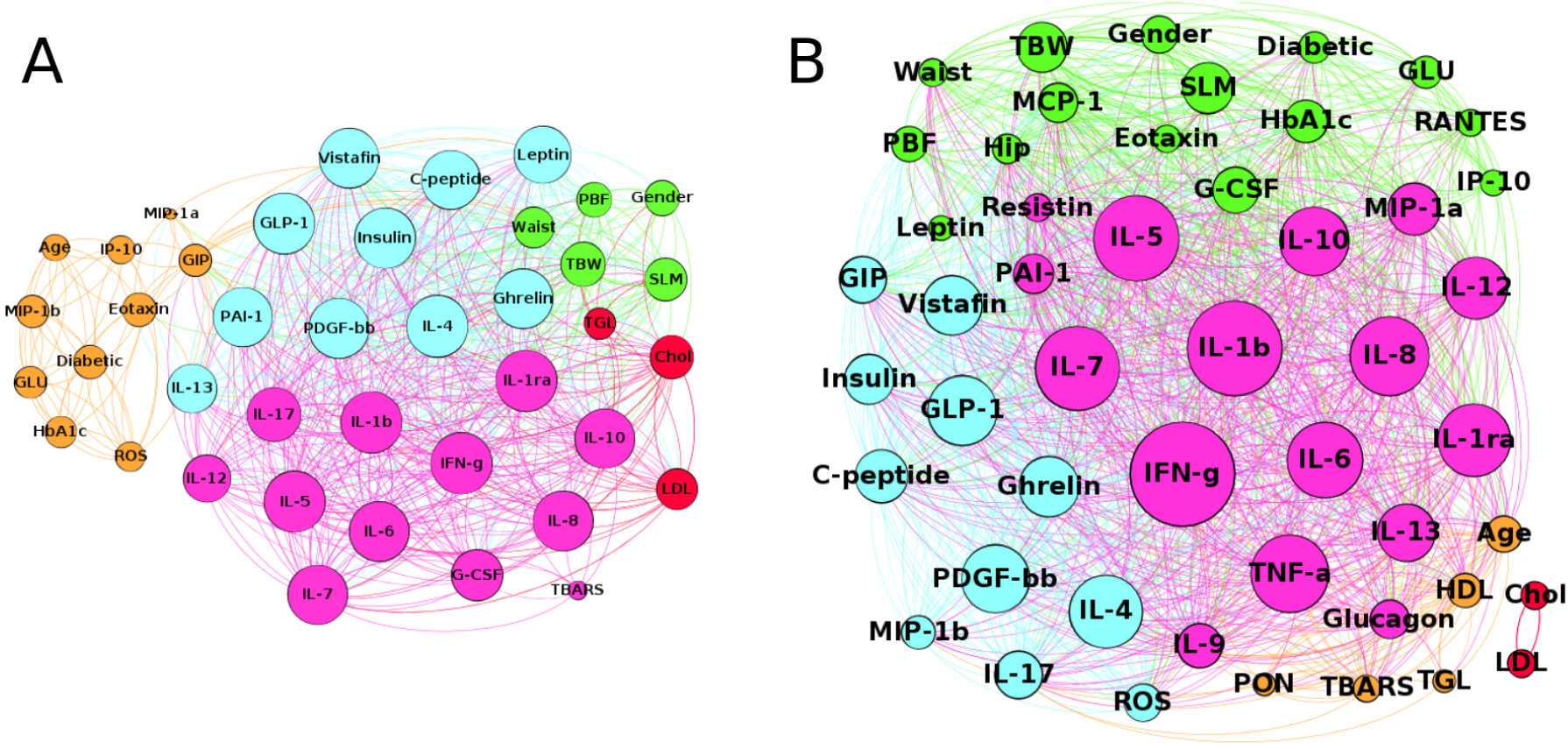
Mutual dependence networks for *G_lean_* and *G_obese_.* Dependence network of characteristics for lean cases. (A) and obese cases (B).

In the case of *G_lean_* there are two main giant connected components corresponding to inflammatory markers (IL*) and metabolic features respectively. There is also presence of two small and compact communities, one corresponding to clinical features like TGL, Chol and LDL while the other corresponds to cluster of physical features like Waist, PBF, TBW, Gender and SLM. A mixed cluster (orange colored) also exists in *G_lean_* whose size and density is more in comparison to the mixed cluster in *G_obese_.* Further, it is apparent from Figure 2A and Figure 2B that there is a strong mutual dependence among the biochemical features resulting in bigger nodes which is proportional to the degree of these variables in the corresponding network.

We observe in *G_obese_* that there is one large community composed primarily of inflammatory markers like IL*, another large community made up of mainly physical features like Waist, PBF, Gender, TBW etc. There is another giant cluster in *G_obese_* consisting of metabolic markers like Insulin, Vistafin, C-peptide, Ghrelin etc along with two small groups where one corresponds to clinical traits like Chol and LDL and the other is a mixed cluster.

Next, we applied the Closed-Form technique (see Material and Methods: Network Based Analysis) to generate the differential subnetworks of *G_lean_* and *G_obese_* as shown in Figure 3. We observe four clusters in the differential subnetwork of *G_lean_* (see Figure 3A) where one community primarily consists of biochemical features, one community comprises physical features and one cluster is made up of clinical features like Chol and TGL. Majority of the nodes present in the mixed cluster of *G_lean_* are part of a community in the differential subnetwork of *G_lean_.* However, the mutual dependence between these features has been reduced to small sized nodes as observed in Figure 3A.

**Fig 3.**
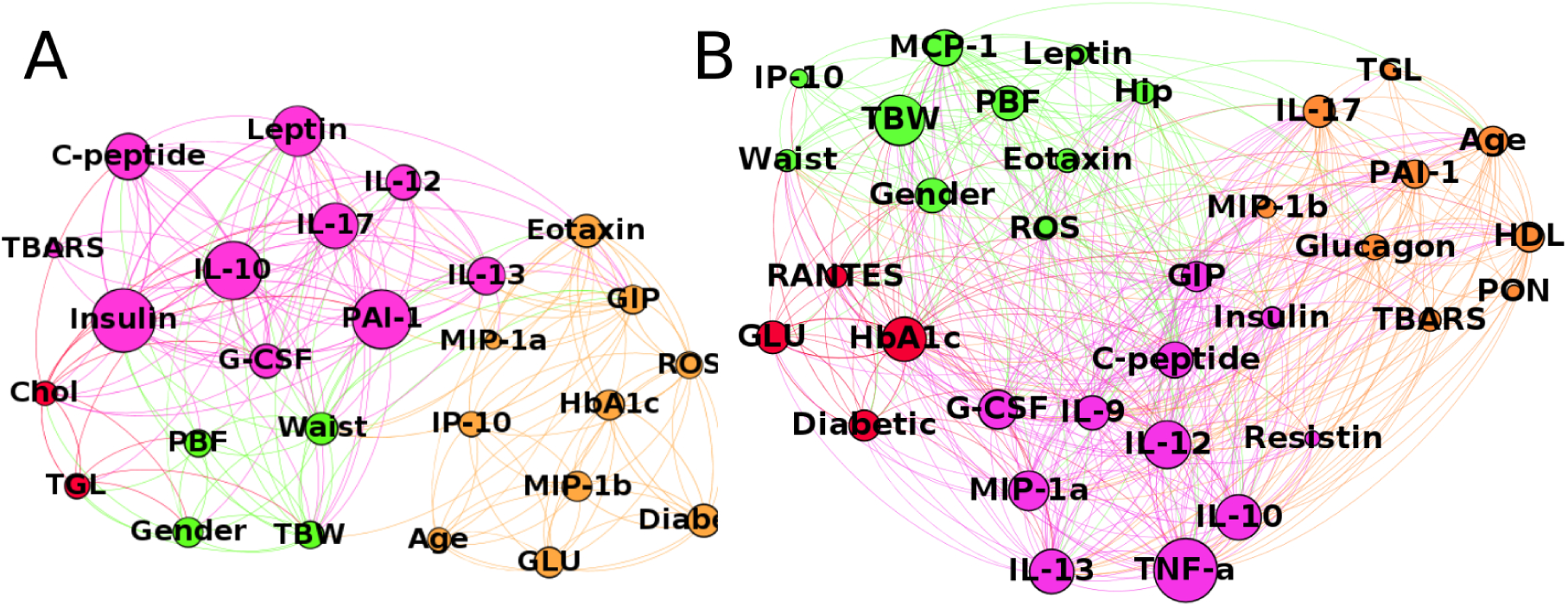
Differential subnetworks for *G_lean_* and *G_obese_.* MI based differential subnetworks of features for lean cases (A) and obese cases (B).

In contrast the differential subnetwork of *G_obes_* (see Figure 3B), though composed of more nodes, is also divided into four communities by the Louvain method. In this network we observe that there exists one community made primarily from physical features and one community composed of mainly biochemical features. Interestingly, we discover one small cluster made up of Glucose (GLU), HbA1c, Diabetic and RANTES. This indicates that the mutual dependence between these features is stronger in *G_obese_* in comparison to *G_lean,_* thereby resulting in a separate community in the differential network of *G_obese._* Several nodes from the mixed cluster of *G_obese_* form a community in the differential subnetwork of *G_obese._* However, the mutual dependence between these characteristics has reduced resulting in smaller size nodes as observed in Figure 3B.

#### Diabetes

In this subsection we report the difference in the effects of the physical, clinical and biochemical features w.r.t. to diabetes by applying the same techniques.

In Figure 4 we illustrate significant MI values of all variable pairs for the dataset 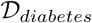 as heat maps. In the non-diabetic subjects, we observe one predominant clusters where the characteristics have low mutual dependence (see Figure 4A) whereas in the diabetic case shown in Figure 4B we see four clusters with relatively higher mutual dependence between the variables within the communities. Next, we applied the same procedure as in the previous subsection to highlight the intricate differences between the non-diabetic and diabetic cases.

**Fig 4.**
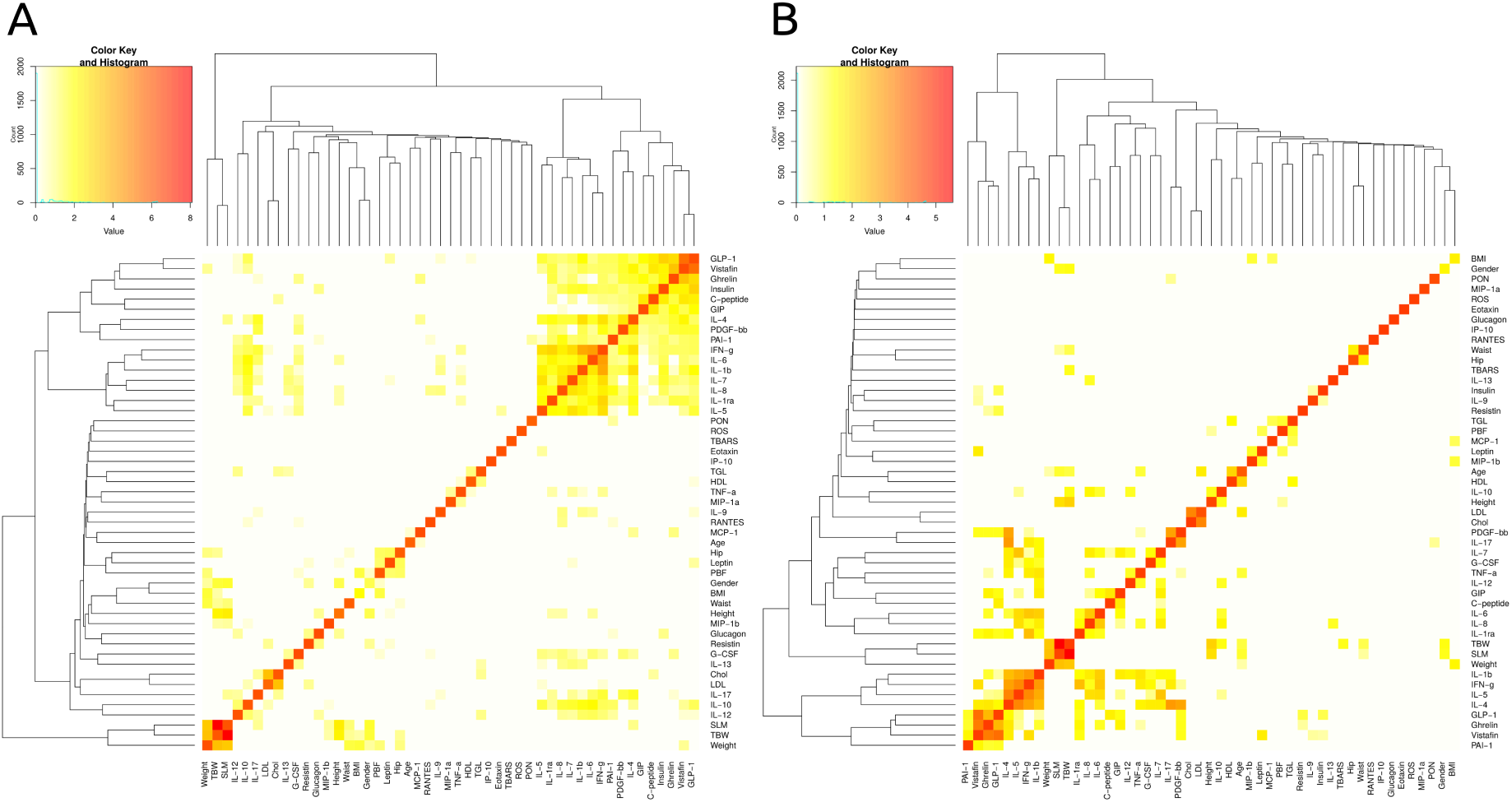
Mutual information heat map for the 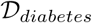 data set. MI based heat map of variables representing non-diabetic cases (A) and diabetic cases (B).

In Figure 5 we represent the mutual dependance networks for non-diabetic *G_control_* (Figure 5A) and diabetic *G_diabetes_* (Figure 5B) subjects. The *G_control_* network consists of 46 nodes with 1348 edges whereas the *G_diabetes_* network is composed of 42 nodes with 682 edges. The *G_control_* network is split into four communities including one corresponding to physical, one clinical, one metabolic and one inflammatory features. It is readily evident from Figure 5A that the nodes have high degree indicating strong mutual dependence.

**Fig 5.**
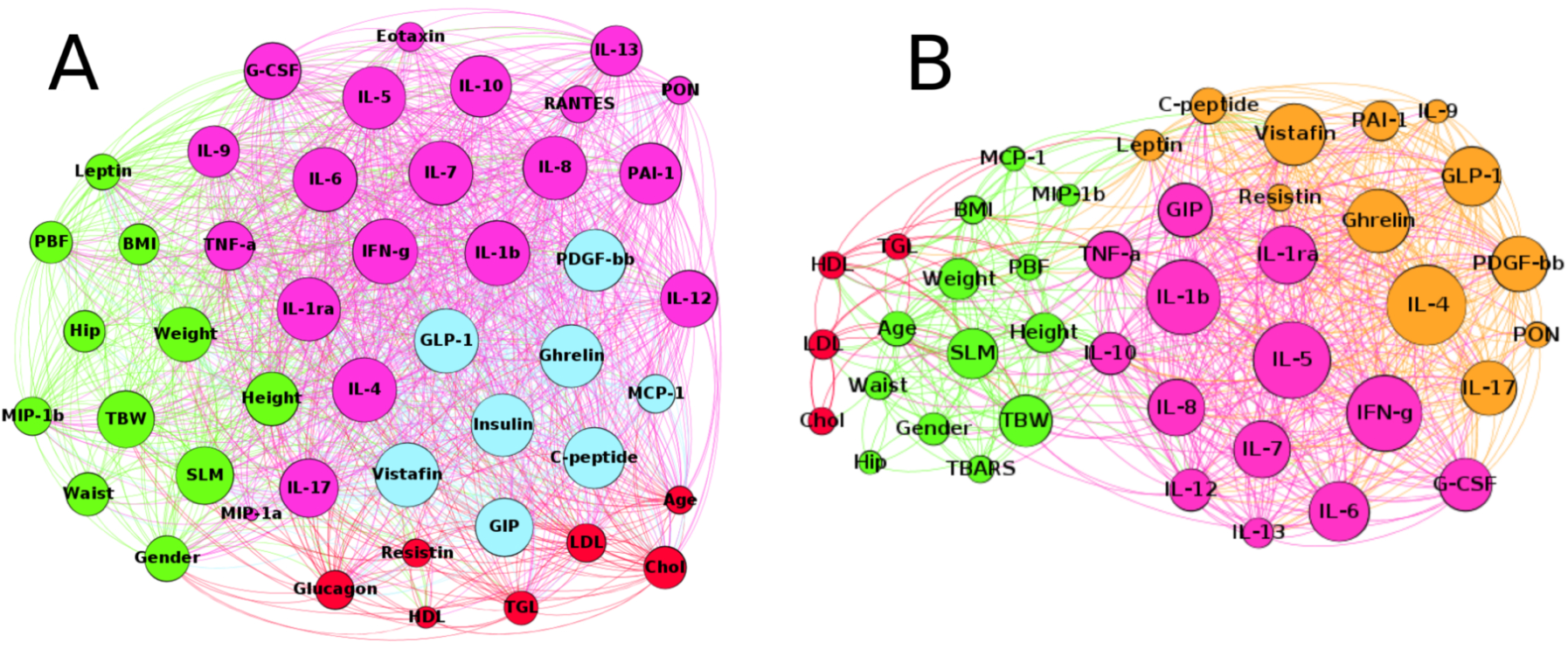
Mutual dependance networks for *G_control_* and *G_diabetes_.* Dependence network of characteristics for non-diabetic cases (A) and diabetic cases (B).

In the *G_diabetes_* network (see Figure 5B) we detect the presence of four communities where one cluster comprises only of clinical features Chol, TGL, HDL and LDL. The are two clusters corresponding to biochemical variables where one is mainly composed of inflammatory features and the second consists of metabolic characteristics. The fourth community is composed primarily from physical features like Age, Weight, Waist, BMI, SLM, Height etc. Interestingly, we noticed that the number of edges, i.e. the mutual dependence between the nodes, is much smaller than in the *G_control_* network.

We applied the Closed-Form method to generate the differential subnetworks for *G_control_* and *G_diabetes_* illustrated in Figure 6. In the control case we detect three coherent communities where one corresponds to biochemical, one to physical and one to clinical features. There is another mixed cluster consisting of several physical and metabolic features. We observe from Figure 6A that the biochemical features retain strong mutual dependence in the case of non-diabetic subjects with a marker like Insulin having a very high mutual dependence with other biochemical traits.

**Fig 6.**
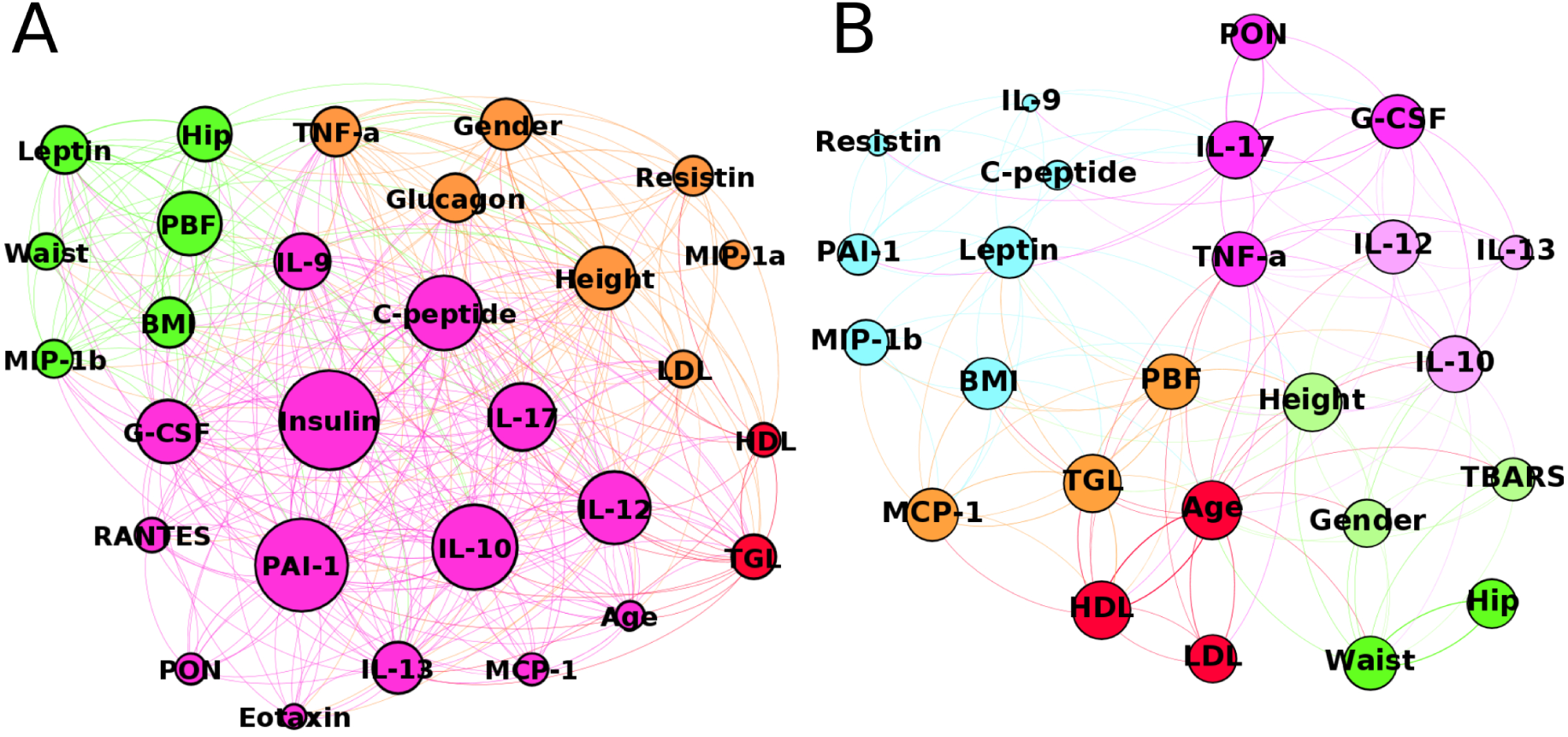
Differential subnetworks for *G_control_* and *G_diabetes_.* MI based differential subnetworks of features for non-diabetic cases (A) and diabetic cases (B).

However, in the differential subnetwork of *G_diabetes_* we observe seven clusters where two clusters belong to inflammatory markers, one big community is made up of metabolic features, two small clusters correspond to physical features and one small community of clinical characteristics. There is also a presence of mixed cluster in the differential subnetwork of *G_diabetes_.* An interesting observation is that Insulin is not present in the community of metabolic markers indicating that in diabetic patients Insulin looses its mutual dependence with other metabolic features.

Apparently, the differential subnetwork of *G_diabetes_* has far fewer edges in comparison to the subnetwork of *G_control_* which indicates that each individual characteristic in the diabetic case is dependent on fewer features than in the control.

## Discussion

In this study, we successfully applied state-of-theart statistical and network analysis techniques on Kuwaiti expression profile data of human subjects with and without T2D. First, we inferred highdimensional models that provide strengths of physical, clinical and biochemical features w.r.t. to lean and obese as well as diabetic and non-diabetic cases. In particular, we used the regularisation methods *elastic net, hdi* and *glinternet.*

We found that PBF and TBW are significantly associate with BMI. This result confirms that waist circumference explains obesity-related risk [46]. Thus, for a given PBF and TBW values, obese and normalweight persons have comparable health risks. However, the other markers such as SLM, HDL, MIP, ROS and RANTES are interesting to investigate especially the latter as it can be a promising therapeutic target for the reduction of NAFLD and NASH (NAFLD: excessive fat accumulation in the form of triglycerides in the liver and has become the most common cause of chronic liver disease in wealthy countries) as was confirmed by [47].

On the other hand, when we used *elastic net* we showed that Diabetes is associated with a significant increase in thiobarbituric acid reactive substances (TBARS) which are considered as an index of endogenous lipid peroxidation as it is explained by [48]. When we used *glinternet,* TBARS was shown to be a marker with the highest coefficient along with thirteen other interactions including those involving Eotaxin and other inflammatory markers. Some of these markers have angiogenic properties, i.e., IL-13, IL-9, while others also contribute to leukostasis and interstitial inflammation, i.e., ROS and the chemokine MIP as explained in [48]. Therefore, eotaxin and co-varying inflammatory markers may be part of a complex pathway resulting in glomerulosclerosis and interstitial fibrosis for patients with T2D as seen in advanced chronic kidney disease [49].

We successfully inferred high-dimensional models that provide effect strengths of physical, clinical and biochemical features w.r.t. lean and obese as well as diabetic and non-diabetic cases. The algorithms work very well as they do not only infer univariate effects of physical, clinical, inflammatory and metabolic markers but also provide pairwise effects via interaction between the variables.

Furthermore, from the mutual dependence networks we observe that the mutual dependence between pairwise features dramatically changes with the phenotype cases. This is reflected in the case of obesity where *G_lean_* is much sparser (has fewer connections) in comparison to *G_obese,_* thereby indicating less dependence of markers on each other. Similarly, in case of diabetes, *G_diabetes_* is much sparser in comparison to *G_control._* A significant observation is that Insulin is not even present in *G_diabetes_* indicating that for diabetic patients Insulin looses its mutual dependence with other metabolic markers as observed in *G_control_.* Another interesting observation is that HbA1c, Glucose (GLU), Diabetic and RANTES form a well-segregated community in the differential sub-network of *G_obese_* whereas they are part of a mixed community in case of differential sub-network of *G_lean._* This indicates that the mutual dependence between these variables is much stronger in the differential sub-network of *G_obese_* in comparison to that of *G_lean_.*

## Conclusion

This case study has several strengths. We used clinically relevant data using human samples. We also used robust statistical tools to analyse our data and established networks based on cross talk between different variables. Our result show that diabetes was associated with a significant increase in thiobarbituric acid reactive substances (TBARS) which are considered as an index of endogenous lipid peroxidation and two inflammatory markers MIP-1 and RANTES. Furthermore, we obtained 13 pairwise effects from *glinternet.* The pairwise effects include pairs from and within physical, clinical and biochemical features, in particular metabolic, inflammatory, and oxidative stress markers. This result confirms for the first time that factors of oxidative stress such as MIP-1 and RANTES participate in the pathogenesis of many diseases such as diabetes and obesity that afflict millions of human subjects. Our results show that markers such as RANTES is interesting to investigate as it can be a promising therapeutic target for the reduction of NAFLD and NASH (NAFLD: excessive fat accumulation in the form of triglycerides in the liver and has become the most common cause of chronic liver disease in wealthy countries).

We would like to point out that the current dataset is relatively small. Nevertheless, the applied techniques provided fairly impressive results. In future, we are looking forward to apply these techniques on larger clinical datasets and team up with experimentalists to verify our findings. Our aim is to encourage researchers in the field to use these techniques for analysis and identification of potential bio-markers from large scale diabetes or obesity data.

## Authorship

Raghvendra Mall performed the network based analysis and provided content for the manuscript. Reda Rawi performed the statistical tests and wrote majority of the manuscript. Ehsan Ullah performed the baseline statistical analysis, generated baseline tables and helped with writing the manuscript. Khalid Kunji generated the figures corresponding to the networks and helped with writing the manuscript. Abdelkrim Khadir, Ali Tiss and Jehad Abubaker collected, cleaned and provided the data in a form on which statistical analysis could be performed. Mohammed Dehbi helped with the biological validity of found traits and provided content for discussion. Halima Bensmail conceived the case study, formulated the objectives of this study, worked on the discussion and helped with validation of the significant clinical markers through thorough literature review.

### Funding

The authors received no specific funding for this work.

### Ethical Approval

The article adheres to principles expressed in Declaration of Helsinki and the ethics committee that approved the study is the Review Board of Dasman Diabetes Institute.

